# Hypoconnectivity between anterior insula and amygdala in neonates with familial history of autism

**DOI:** 10.1101/2022.02.24.481850

**Authors:** Dustin Scheinost, Joseph Chang, Cheryl Lacadie, Emma Brennan-Wydra, Rachel Foster, Alexandra Boxberger, Suzanne Macari, Angelina Vernetti, R. Todd Constable, Laura R. Ment, Katarzyna Chawarska

## Abstract

**Background:** Altered resting state functional connectivity (FC) involving the anterior insula (aINS), a key node in the salience network, has been reported consistently in autism.

**Method:** Here we examined, for the first time, FC between the aINS and the whole brain in a sample of full-term, postmenstrual age (PMA) matched neonates (mean 44.0 weeks, SD=1.5) who due to family history have high likelihood (HL) for developing autism (n=12) and in controls (n=41) without family history of autism (low likelihood, LL). Behaviors associated with autism were evaluated between 12 and 18 months (M=17.3 months, SD=2.5) in a subsample (25/53) of participants using the First Year Inventory (FYI).

**Results:** Compared to LL controls, HL neonates showed hypoconnectivity between left aINS and left amygdala. Lower connectivity between the two nodes was associated with higher FYI risk scores in the social domain (r(25) = −.561, p=.003) and this association remained robust when maternal mental health factors were considered. Considering that a subsample of LL participants (n=14/41) underwent brain imaging during the fetal period at PMA 31 and 34 weeks, in an exploratory analysis, we evaluated prospectively development of the LaINS-Lamy connectivity and found that the two areas strongly coactivate throughout the third trimester of pregnancy.

**Conclusions:** The study identifies left lateralized anterior insula - amygdala connectivity as a potential target of further investigation into neural circuitry that enhances likelihood of future onset of social behaviors associated with autism during neonatal and potentially prenatal periods.

## Introduction

Autism spectrum disorder (referred to henceforth as ‘autism’) is an early-onset neurodevelopmental disorder characterized by atypical social interaction patterns and presence of restrictive interests and repetitive behaviors.^1^ Genetic factors involved in the etiology of autism are thought to affect development of structural and functional brain networks, which in turn contribute to the development of behaviors constituting core autism symptoms. A functional network heavily implicated in autism is the salience network, consisting of anterior insula (aINS) and anterior cingulate cortex (ACC).^2–5^ The aINS is a major afferent hub that receives extensive viscero-autonomic feedback, while the ACC functions as an efferent hub responsible for generating relevant visceral, autonomic, behavioral, and cognitive responses.^6,7^ The salience network links other cortical networks with limbic and autonomic systems, and is involved in detection of and orienting to salient external and internal stimuli, including those that are social in nature.^7^

There is extensive evidence based on studies with preschoolers, school-age children, and adolescents with autism of reduced communication between aINS and other cortical nodes including those comprising the default mode (DMN) and the frontoparietal (FPN) networks, as well as the amygdala.^4,8–13^ This reduced functional connectivity (FC) may result in difficulties in transfer and integration of information across these networks, which then could contribute to the impaired processing of social signals in autism. However, given that autism onsets in early childhood and that most FC studies are conducted in older children, it is not clear to what extent the observed differences precede emergence of symptoms of autism or emerge secondary to the way children with autism experience and interact with their social and nonsocial environment.

The aINS is one of the first cortices to differentiate and develop in utero,^14^ providing structure for functional specialization at the time of birth.^15^ The connections between aINS and ACC nodes undergo dramatic growth in the first months of life, with the network reaching near adult-like characteristics by 12 months.^16,17^ Similarly, the connectivity between the aINS node and the nodes in other cortical networks increases rapidly over the course of the first year, including those in the DMN and FPN networks as well as with the amygdala, indicating increasing synchronization and integration between the networks.^16,18,19^ Given the relatively late maturation of the higher order networks, the insula showing functional specialization at birth may indicate its special importance for early survival.^15^ Considering that the anterior insula receives and integrates exteroceptive and interoceptive input, aids in the ongoing maintenance of homeostasis, and maintains rich connections with the subcortical structures involved in emotional processing and valuation, it has been hypothesized that aINS plays a key role in development of social behaviors and motivation in neonates.^20^

Extant, albeit still very limited, evidence suggests that increased genetic risk factors for autism exert their effect on development of FC shortly after birth. Younger siblings of children with autism, who due to genetic factors have a higher likelihood (HL) for developing autism than the general population, show alterations of FC patterns during the first two post-natal months. These alterations include increased regional homogeneity (ReHo) in the fusiform gyrus and parietal cortex nodes,^21^ increased within-network connectivity in the DMN, ^22^ and decreased connectivity between right aINS and ACC and other higher order cortical regions compared to neonates without genetic risk factors for autism (i.e., low likelihood (LL) for developing autism).^23^

Here, we examined differences in the aINS seed-based FC in HL and LL neonates. Based on prior work with older children with autism, we hypothesized that HL neonates would exhibit reduced aINS connectivity with other cortical areas compared to the LL controls. We further hypothesized that the reduced aINS connectivity would be associated with elevated risk scores for autism-related behaviors measured later in infancy. Considering reports regarding links between maternal mental health and developing connectome, ^24–30^ as well as infant social and emotional development,^31–33^ indices of maternal stress, anxiety, and depression were controlled for in the analyses.

## Methods

### Participants

The participants included 63 (14 HL, 49 LL) neonates. The neonatal scans were completed between April 11, 2018 and December 14, 2021 as part of a prospective study of FC development in autism. Parents provided written informed consent to participate in the study prior to data acquisition. The study was conducted in accordance with the Strengthening the Reporting of Observational Studies in Epidemiology (STROBE) reporting guideline for cohort studies. Inclusion criteria were singleton pregnancy, term birth, and appropriate for gestational age birthweight. HL neonates had to have an older sibling with autism diagnosis confirmed through record review. LL neonates were excluded if they had a first or second degree relative with autism. Additional exclusions for all participants were: 1) congenital infections; 2) nonfebrile seizure disorder; 3) hearing loss; 4) visual impairment; 5) known genetic abnormality; 6) prenatal exposure to illicit drugs; 7) major psychotic disorder in a first degree relative; and 8) contraindications to MRI including non-removable metal medical implants (e.g., patent ductus arteriosus clip). Ten out of 63 (15.87%) neonates did not contribute valid MRI data due poor registration (n=4), motion artefacts (n=2), and lack of functional runs (n=4).

The sample retained for the analysis included 53 neonates (12 HL and 41 LL) scanned between 41.57 and 49.85 postmenstrual (PMA) weeks (see **Table 1**). Males constituted 83% of the HL and 46% of the LL sample (χ^2^(1) = 5.127, p=.024) (see **Table 1**). 33% of the HL sample and 29% of the LL sample reported a racial background other than white and 25% of the HL sample and 15% of the LL sample were ethnically Hispanic or Latino. 83% of HL mothers and 97% of LL mothers held a bachelor’s degree or higher. The two groups did not differ in maternal age, race, ethnicity, or maternal education (all p>.080) at the time of scan (see **Table 1**). All neonates have been participating in an ongoing prospective longitudinal study and their social development is assessed periodically in the second year of life, with diagnostic outcome measures expected to be completed at the age of 24 months. At 24 months, the participants undergo a clinical evaluation using the Mullen Scales of Early Learning (MSEL) capturing verbal and nonverbal developmental skills, the Autism Diagnostic Observation Schedule, Second Edition (ADOS-2) Toddler Module quantifying symptoms of autism, and the Vineland Adaptive Behavior Scales, Third Edition (VABS-3) capturing adaptive skill level. Due to genetic liability, approximately 20% of HL participants are expected to develop autism, whereas another 30-40% are expected to demonstrate delays and atypical features often referred to as broader autism phenotype (BAP) by the time they reach 2 years of age.^34–36^

**Table 1.** Sample characterization

|  | N | HL | N | LL | p-value |
| --- | --- | --- | --- | --- | --- |
| **Demographics** |  |  |  |  |  |
| Male (%) | 12 | 10 (83.3) | 41 | 19 (46.3) | .024 |
| Hispanic/Latino (%) | 12 | 3 (25.0) | 41 | 6 (14.6) | .400 |
| Race (% Non-White) | 12 | 4 (33.3) | 41 | 12 (29.3) | .787 |
| Maternal Age at Scan | 12 | 34.35 (4.02) | 41 | 33.74 (3.87) | .636 |
| Maternal education (% College) | 12 | 10 (83.3) | 37 | 36 (97.3) | .080 |
| Maternal EPDS score | 12 | 5.83 (4.39) | 41 | 4.61 (3.34) | .304 |
| Maternal STAI Trait score | 12 | 34.08 (9.68) | 41 | 32.12 (7.95) | .478 |
| Maternal PSS score | 12 | 18.33 (7.91) | 41 | 17.71 (3.40) | .779 |
| **Neonatal brain imaging** |  |  |  |  |  |
| PMA at scan (weeks) | 12 | 43.94 (1.18) | 41 | 44.08 (1.60) | .780 |
| Valid Scan Time | 12 | 11.62 (0.61) | 41 | 11.38 (1.88) | .659 |
| Frame-to-frame displacement | 12 | 0.04 (0.03) | 41 | 0.03 (0.02) | .211 |
| **Behavioral follow-up** |  |  |  |  |  |
| Age (months) | 9 | 17.26 (2.64) | 16 | 17.28 (2.58) | .986 |
| FYI Social Communication score | 9 | 8.63 (7.96) | 16 | 3.00 (4.05) | .027 |
| FYI Sensory Regulation score | 9 | 6.53 (6.12) | 16 | 3.89 (5.96) | .304 |

### Maternal mental health

Maternal perinatal mental health was quantified using the Edinburgh Postnatal Depression Scale (EPDS),^37^ a 10-item measure of perinatal depression yielding possible scores ranging from 0 to 30; the State-Trait Anxiety Inventory (STAI),^38^ a 40-item measure of both state and trait anxiety total scores each ranging from 20 to 80; and the Perceived Stress Scale (PSS-14),^39^ a 14-item measure of perceived stress with scores ranging from 0 to 56. For all three measures, higher scores indicate the presence of more symptoms.

### Autism-related social and regulatory behaviors

Behaviors considered risk factors for autism were assessed using the First Year Inventory 2.0 (FYI) parent questionnaire.^40,41^ The FYI measure is sensitive to autism-specific behaviors in the general population and amongst younger siblings of children with autism^42–46^ and shows positive correlations with concurrent direct measures of autism symptom severity captured by the ADOS-2 Toddler Module^47^ and the Autism Observation Scale for Infants (AOSI).^48^ Unlike the ADOS-2, the FYI was designed to capture autism traits in the general population, and thus, is well-suited for correlational studies in neurodiverse groups. The FYI consists of Social Communication (28 items) and Sensory-Regulatory (24 items) domains. Risk scores were generated based on assigned risk points derived from the normative sample.^40^ Each construct score ranges from 0 to 50 on a semi-logarithmic scale, with higher scores indicating more symptoms.

### Imaging parameters

Participants were scanned without sedation during natural sleep using the feed-and-wrap protocol.^49^ Infants were fed, bundled with multiple levels of ear protection, and immobilized in an MRI-safe vacuum swaddle. Heart rate and O_2_ saturation were continuously monitored during all scans. The scans were performed using a 3 Tesla Siemens (Erlangen, Germany) Prisma MR system with a 32-channel parallel receiver head coil. Functional runs were acquired using a multiband T2*-sensitive gradient-recalled, single-shot echo-planar imaging pulse sequence (TR = 1 s, TE = 31 ms, FoV = 185 mm, flip angle 62°, multiband = 4, matrix size 92 × 92). Each volume consisted of 60 slices parallel to the bi-commissural plane (slice thickness 2 mm, no gap). We collected 4-5 functional runs, each comprised of 360 volumes. Each neonate had on average 11.6 min (SD = 1.34) of usable functional data with an average frame-to-frame displacement of 0.03 (SD=0.02, Min: 0.01, Max: 0.10). High-resolution T1-and T2-weighted 3D anatomical scans were acquired using an MPRAGE sequence (TR=2400 ms, TE=1.18 ms, flip angle=8°, thickness=1 mm, in-plane resolution=1 mm x 1 mm, matrix size=256 × 256) and a SPACE sequence (TR=3200 ms, TE=449 ms, thickness=1 mm, in-plane resolution=1 mm x 1 mm, matrix size=256 × 256).

### Common Space Registration

Functional images were warped to a custom infant template using a series of linear and nonlinear registrations. These registrations were calculated independently and combined into a single transform. This allows the participant images to be transformed to common space with only one transformation, reducing interpolation error. Functional images were linearly registered to the anatomical image, which was nonlinearly registered to the infant template using a previously validated algorithm.^50^

### Connectivity processing

Functional data were processed using a previously validated pipeline.^49^ Functional images were slice-time and motion corrected using SPM8. Next, images were iteratively smoothed until the smoothness of any image had a full-width half maximum of approximately 6 mm using AFNI’s 3dBlurToFWHM. This iterative smoothing reduces motion-related confounds.^51^ All further analyses were performed using BioImage Suite^52^ unless otherwise specified. Several covariates of no interest were regressed from the data including linear and quadratic drifts, mean cerebral-spinal-fluid (CSF) signal, mean white-matter signal, and mean gray matter signal. For additional control of possible motion-related confounds, a 24-parameter motion model (including six rigid-body motion parameters, six temporal derivatives, and these terms squared) was regressed from the data. The data were temporally smoothed with a Gaussian filter (approximate cutoff frequency=0.12Hz). A canonical gray matter mask defined in common space was applied to the data, so only voxels in the gray matter were used in further calculations.

### Seed connectivity

We assessed whole brain seed connectivity from seeds in the right and left aINS, as shown in **Supplemental Figure S1**. Seeds were manually defined on the reference brain. The time course of the reference region in each participant was then computed as the average time course across all voxels in the seed region. This time course was correlated with the time course for every other voxel in gray matter to create a map of r-values, reflecting seed-to-whole-brain connectivity. These r-values were transformed to z-values using Fisher’s transform yielding one map for each seed and representing the strength of correlation with the seed for each participant.

### Head Motion

Since head motion can potentially confound FC, we included several steps to ensure adequate control of motion confounds. For neonates, the mean frame-to-frame displacement was calculated for each run for every individual and runs with a mean frame-to-frame displacement greater than 0.10 mm were removed from further analysis. Additionally, iterative smoothing, regression of 24 motion parameters (6 rigid-body parameters, 6 temporal derivatives of these parameters, and these 12 parameters squared), and regression of the global signal were used in the neonatal data.

### Exploratory fetal analysis

A subsample of the LL neonates (14/41) contributed brain imaging data during the fetal period at 31 (n=12) and 34 (n=14) PMA weeks; 2 were scanned at 2 time points, 12 were scanned at all 3 time points. In this exploratory analysis, we leveraged the longitudinal FC MRI data collected in the LL group to evaluate normative fetal-to-neonatal development of the connections identified as atypical in the HR sample. All MRIs were performed in the natural, unmedicated state. Fetuses were studied using repeat MRI protocols completed in <60 minutes using a 3 Tesla Siemens (Erlangen, Germany) Prisma MR system and a flexible, lightweight (∼1lb), cardiac 32-channel body coil. Five functional runs were acquired (TR=2000 ms, TE=30 ms, FoV=352×400mm, flip angle 90°, matrix size 88×100, SAR<0.4, slice thickness 3mm, Bandwidth=2940 Hz/pixel, 32 slices). Each of the 5 functional runs were comprised of 150 volumes (5 minutes). All participants had at least 5 minutes of usable data, with an average of 16.7 min (SD = 7.18) for the 31-week scan and 20.4 min (SD = 5.36) for the 34-week scan.

Functional data were processed using validated fetal fMRI pipelines.^53,54^ Functional data were corrected for motion using a two-pass registration approach optimized for fetuses to correct for large and small head movements.^53^ Outlying frames were censored for data quality based on signal-to-noise (SNR) ratio within the fetal brain, the final weighted correlation value from optimization, and the frame-to-frame motion between adjacent frames. These frames were defined as frame with SNR, registration quality, or motion greater/less than one standard deviation above/below the mean values over all runs. As outlined in Thomason et al.,^55^ several covariates of no interest were regressed from the data including linear and quadratic drifts, 6 motion parameters, mean cerebral-spinal-fluid (CSF) signal, mean white-matter signal, and mean gray matter signal. The data were temporally smoothed with a zero mean unit variance Gaussian filter (approximate cutoff frequency = 0.12 Hz).

Next, the left aINS and amygdala seeds from infant space were warped to fetal fMRI space for analyses. First, the mean functional image from the motion corrected fMRI data was registered to an age-appropriate template (i.e., 31 weeks or 34 weeks gestation)^56^ using a low-resolution non-linear registration. The fetal templates were non-linearly registered to a custom infant template using the same algorithm as above.

### Statistical analysis

Imaging data were analyzed using voxel-wise two-sample t-tests. The significance threshold for imaging results was set at p < 0.05, with all maps corrected for multiple statistical comparisons across gray matter using cluster-level correction estimated via AFNI’s 3dClustSim (version 16.3.05) with 10,000 iterations, an initial cluster forming threshold of p=0.001, the gray matter mask applied in preprocessing, and a mixed-model spatial autocorrelation function (ACF). Parameters for the spatial ACF were estimated from the residuals of the voxel-wise linear models using 3dFWHMx. Subsequently, using an ANOVA, we evaluated the effects of familial likelihood and sex on the left aINS – left amygdala connectivity with PMA and maternal stress, depression, and anxiety indices as covariates. Significant interaction was followed by post-hoc contrasts with Tukey correction for multiple comparisons. Associations between the left aINS – left amygdala connectivity in neonates and later FYI risk scores in social-communication and sensory-regulatory domains were analyzed using Pearson’s r correlation coefficient while controlling for maternal mental health indices. In an exploratory analysis, we examined longitudinal changes in aINS – amygdala connectivity between 31- and 44-weeks PMA in LL participants using linear mixed effects model. The model included random effects to account for multiple longitudinal measurements from the same individual, incorporated nonlinearity by allowing the slope to change at 35 weeks and allowed for heteroscedasticity by using different error variance parameters to describe the first and second fetal measurements (denoted by F1 and F2) and the neonatal measurement (denoted by NN).

## Results

### Preliminary analyses

There were no differences between the HL and LL groups in the amount of valid resting-state functional connectivity data or the average FTF displacement values (**Table 1**). At the time of writing the manuscript, 25/53 of the study participants (9 HL and 16 LL) reached their second year of life when the FYI questionnaire data was collected at the average 17.27 (SD=2.54) months. There were no differences between the groups in the age at FYI administration, but consistent with prior studies, ^44,57^ HL infants exhibited higher scores in the FYI Social Communication but not in the Sensory-Regulatory domain (**Table 1**). There were also no differences between the groups in terms of maternal perinatal stress, depression, or anxiety indices (**Table 1**). Diagnostic outcomes were available for 20/53 of the participants, determined at a mean age of 23.73 (2.62) months; the remaining children have not reached the diagnostic assessment age yet. Amongst the 12 HL participants scanned as neonates, two were diagnosed with autism, three were noted to have BAP features, two were developing typically and five have not completed their diagnostic assessment yet due to age or COVID-19 pandemic restrictions. In the LL group, two participants were noted to have some developmental delays, 11 were developing typically, and 28 have not completed their diagnostic assessment yet.

### Anterior insula seed based functional connectivity

For the LL group, activity in the left aINS correlated with local activity in surrounds of the aINS, the right aINS, bilateral amygdala, bilateral hippocampus, and dACC (**Figure 1A**). Connectivity patterns of the aINS were similar albeit with reduced connectivity to the amygdala and hippocampus (**Figure 1B**). When compared to the LL group, the HL group had significantly weaker connectivity between the left aINS and the left amygdala (p<0.05 corrected; **Figure 1C**). No significant difference between the HL and LL groups were detected using the right aINS seed.

**Figure 1:**
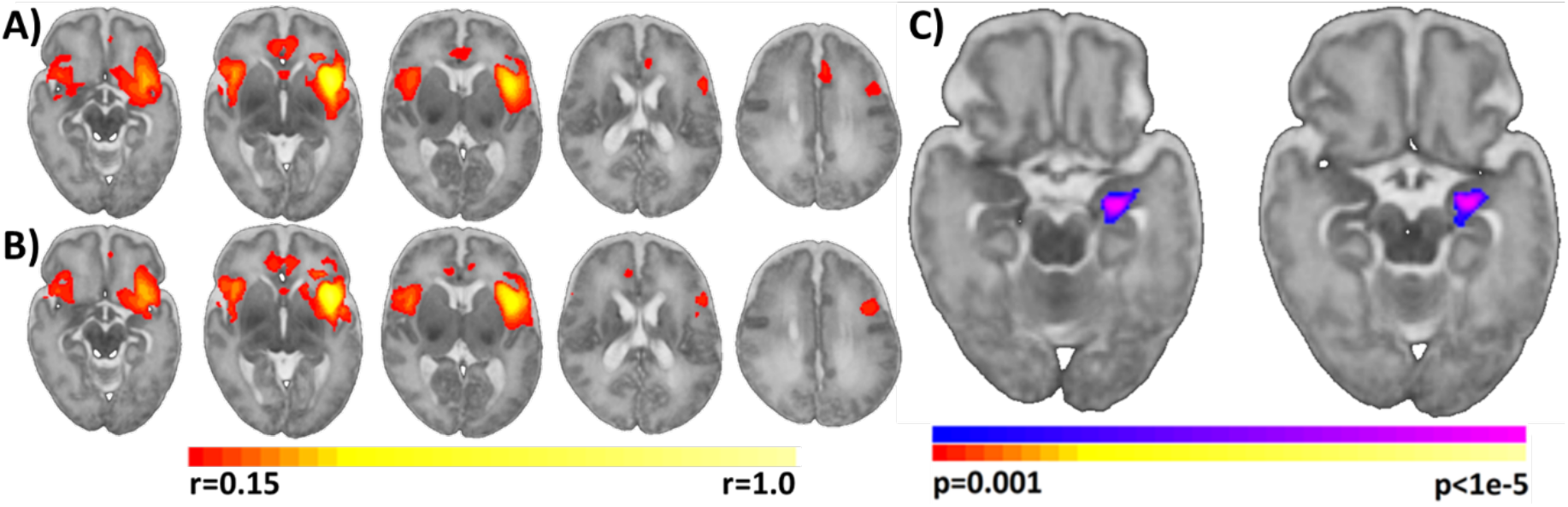
Left anterior insula (aINS) connectivity in neonates with high and low genetic likelihood for developing autism. A) aINS connectivity for the LL group. B) aINS connectivity for the HL group. C). Comparison of aINS connectivity between LL and HL neonates. Overall, the HL group had significant weaker connectivity between the left aINS and the left amygdala compared to the LL group (p<0.05, corrected). Slices are shown in radiological convention.

Considering significant differences in sex ratios in the HL and LL groups, we extracted left aINS – left amygdala connectivity values and conducted a post-hoc group x sex ANOVA, with PMA as a covariate. The ANOVA indicated a significant effect of group, F(1, 48) = 26.68, p<.001, no effect of sex, F (1, 48) = .38, p=.539, and a significant group x sex interaction, F (1, 48) = 4.28, p=.044. The contribution of PMA to the model was negligible, F (1, 48) = .09, p=.761. Post-hoc comparisons with Tukey correction for multiple comparisons indicated that while HL females (M=0.12, SD=0.01) had statistically comparable left aINS – left amygdala connectivity to LL females (M=0.15, SD=0.09) (Cohen’s d = .468, p=.981) and LL males (M=0.15, SD=0.10) (Cohen’s d = .422, p=.972), the HL males (M=-0.04, SD=0.10) showed left aINS – left amygdala hypoconnectivity compared to LL females (Cohen’s d = 2.00, p<.001) and LL males (Cohen’s d = 1.90, p<.0001) (**Figure 2**). Although HL females tended to have higher left aINS – left amygdala connectivity than HL males, the contrast did not survive correction for multiple comparisons (p=.151; uncorrected: p=.036); there were no differences between LL males and LL females (p=.997). While connectivity in the LL group was significantly greater than 0 (M=.15, SD=.0.09), t(40) = 10.26, p<.0001, in the HL group it was not (M=-.01, SD=-0.11), t(11) = −0.39, p=.073.

**Figure 2.**
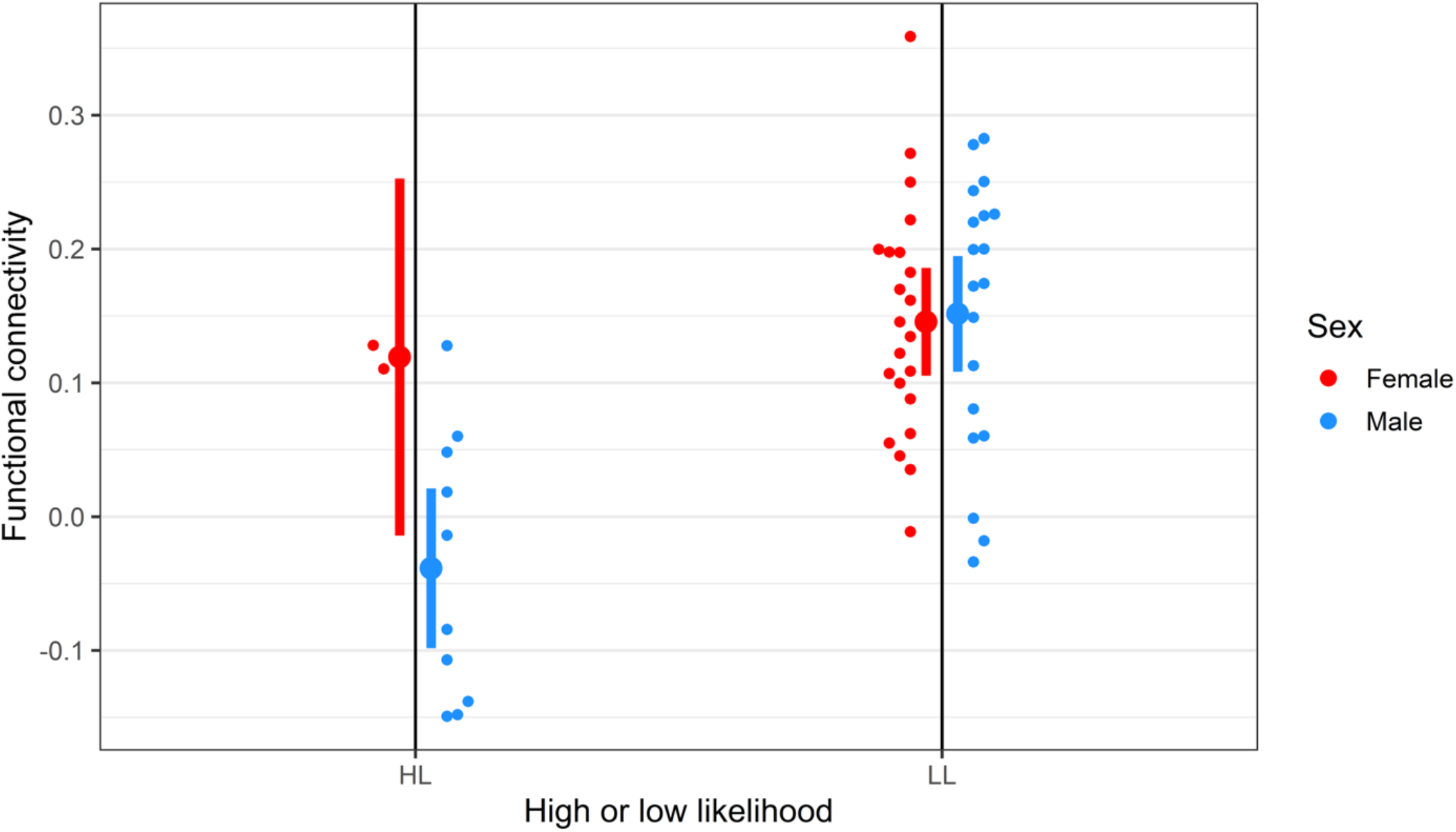
Individual left anterior insula – left amygdala functional connectivity in male (blue) and female (red) neonates with high (HL) and low likelihood (LL) for developing autism. Small dots represent individual data points, large dots represent group averages, and the vertical bars represent 95% confidence intervals.

Given that altered amygdala circuitry in infants is a common sequela of experiencing prenatal maternal distress,^24–30^ we repeated our primary analyses controlling for maternal scores on the EPDS, STAI, and PSS questionnaires. The ANOVA effects remained significant when controlling for the three maternal mental health variables: group: F(1, 46) = 6.23, p=.016), sex: F(1, 46) = 3.57, p=.065. group x sex: F (1, 46) = 4.30, p=.044, with negligible effects of maternal depression (p=.522), trait anxiety (p=.521), and stress (p=.851).

### Associations between functional connectivity and phenotypic features

Subsequently, we computed Pearson’s r correlation between the FYI scores and the left aINS – left amygdala connectivity in the combined neurodiverse sample. The correlation indicated that lower connectivity between left aINS and left amygdala was associated with higher later FYI scores, r(25) = −.561, p=.004 in the Social Communication but not in the Sensory and Regulatory domain, r (25) = −.302, p=.142 (**Figure 3**). The association between aINS - amygdala connectivity and FYI Social communication scores remained robust when maternal EPDS, STAI, and PSS scores were partialled out, r(25) = .-526, p=.012.

**Figure 3.**
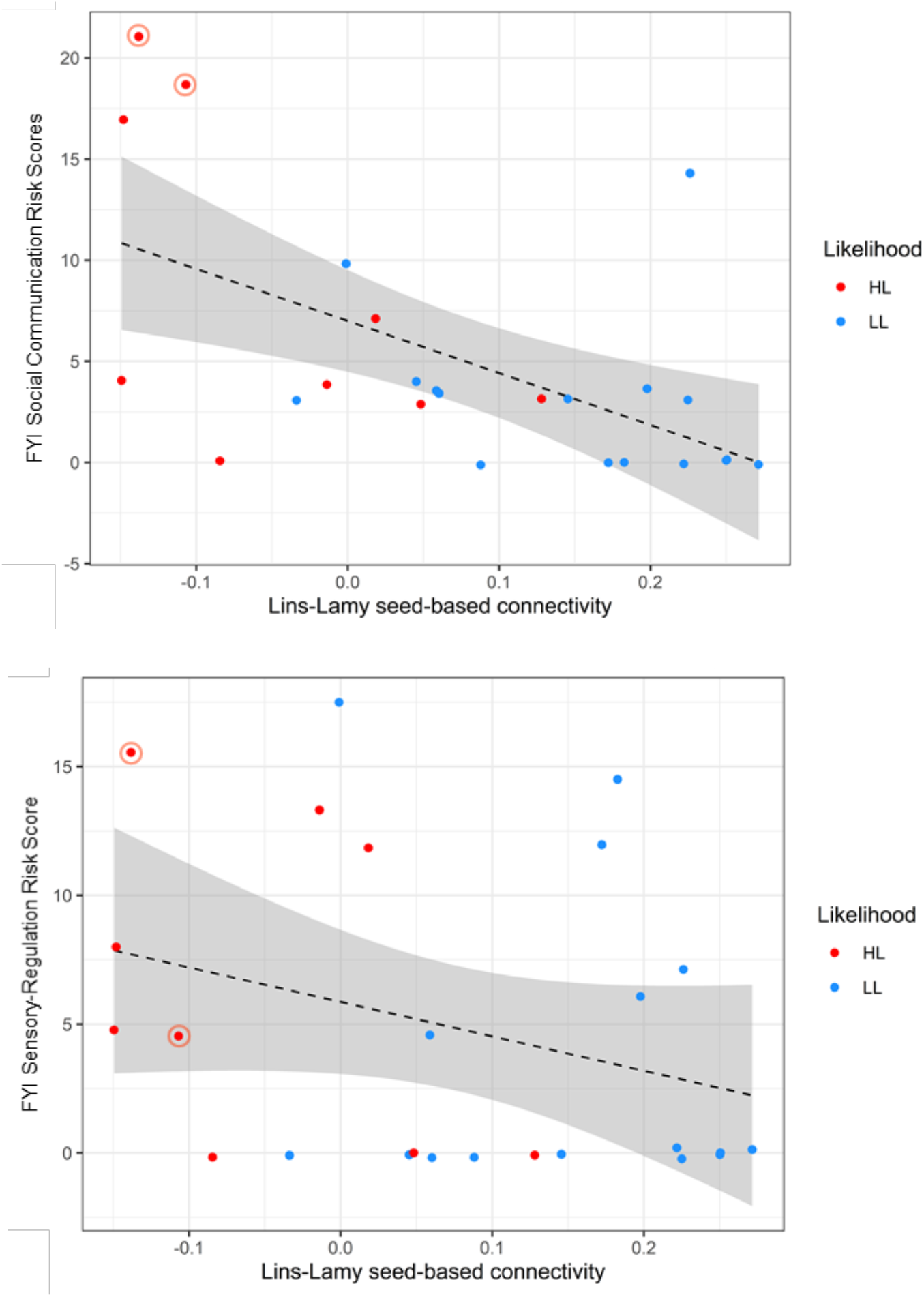
Pearson’s r correlation between left anterior insula – left amygdala connectivity during neonatal period and the First Year Inventory Social Communication (a) and Sensory and Regulatory (b) risk scores collected in the second year of life in HH (red) and LL (blue) infants. Highlighted data points in red represent HL infants later diagnosed with autism. Gray area represents the 95% confidence intervals.

### Development of insula-amygdala edge from fetal to neonatal period in LL only

Finally, we investigated the normative fetal-to-neonatal developmental trajectory of the left aINS – left amygdala connectivity in the LL sample. The data are shown in **Figure 4**, along with the 95% confidence intervals for the mean connectivity at each of the 3 representative ages of 31-, 35-, and 44-weeks PMA. This analysis indicated that connectivity between the two nodes is already greater than 0 at 31 weeks of gestation and does not undergo marked increases or decreases during the prenatal to neonatal transition. Specifically, at each age, the connectivity on this edge was found to be significantly positive, with a mean of .148 at 31 weeks PMA (t(30)=4.90, p<.0001), .136 at 35 weeks (t(30)=5.64, p<.0001), and .122 at 44 weeks (t(30)=9.93, p < .0001) and the linear mixed effects model indicated no effects of PMA, with no significant slope before 35 weeks PMA (mean = −.003/week, p=0.77) and no significant increase in slope at 35 weeks PMA (mean = .0014/week, p=.91).

**Figure 4.**
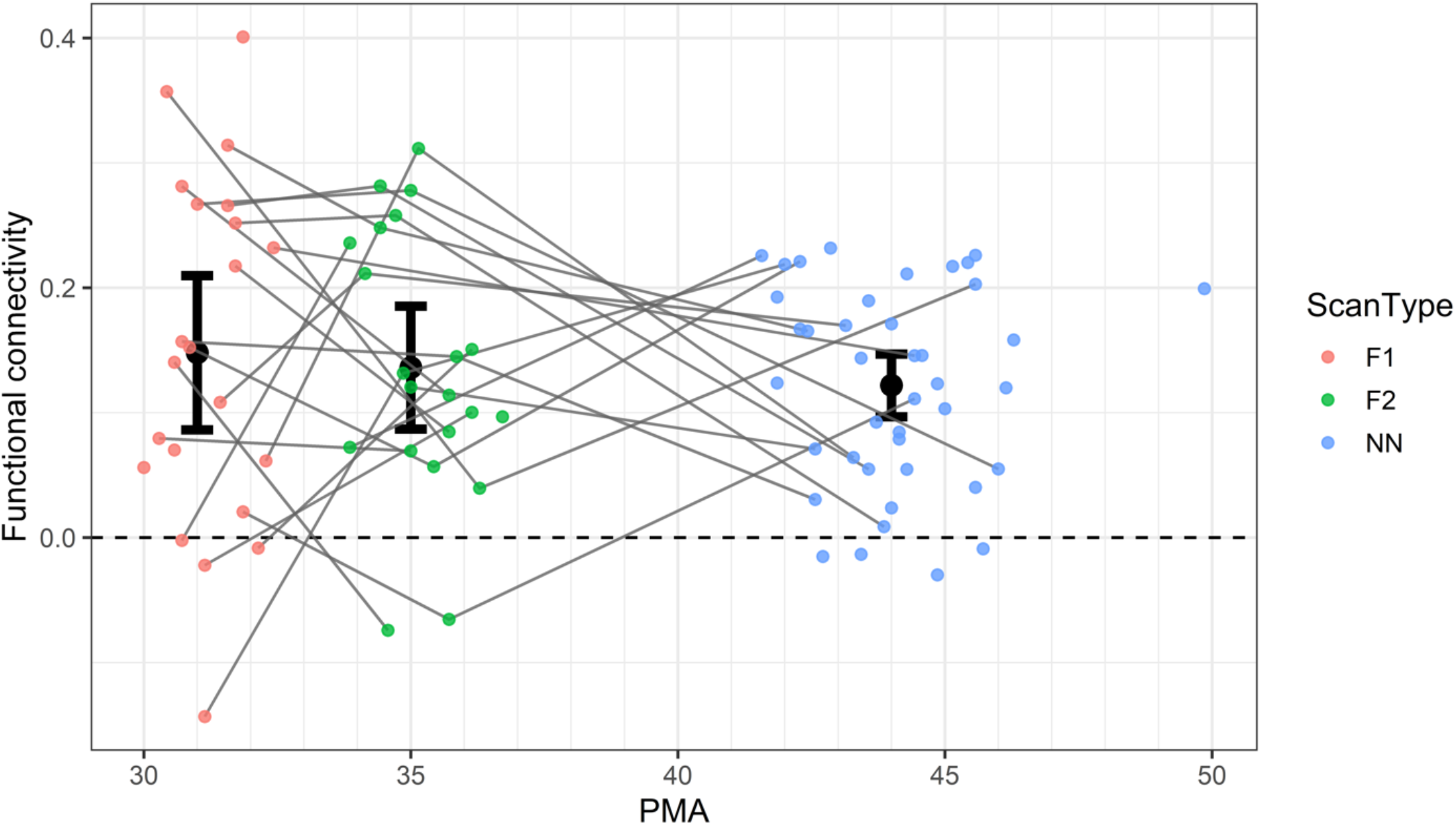
Average (+/− 2 standard errors) left anterior insula – left amygdala functional connectivity in a sample with low likelihood of developing autism at 31 (F1, red)), 35 (F2, green), and 44 (NN, blue) postmenstrual weeks.

## Discussion

Using a prospective longitudinal study design, with data collection from the fetal through toddler period, this study demonstrates left lateralized hypoconnectivity between anterior insula and amygdala in neonates with familial history of autism. Clinical implications of this finding are demonstrated by robust links between lower connectivity between the two nodes and more atypical social communication behaviors recorded 12 to 18 months later. Of note, the observed effects are independent of maternal stress, anxiety, and depressive symptoms, which are known to affect development of the amygdala circuitry^24–30^ as well as social and emotional outcomes in infancy. ^31–33^ The study demonstrates that genetic predisposition for autism manifests in decreased communication between early maturing cortical nodes involved in salience detection. Taken together, these results suggest that anterior insula hypoconnectivity previously reported in older children with autism may be present already during the perinatal period, and that it may first manifest in reduced communication with the amygdala.

Similar to the anterior insula,^14^ the amygdala develops rapidly during the last trimester of pregnancy. At birth, the amygdala shows adult-like positive connectivity with subcortical nodes including the thalamus, caudate, and putamen, which together play a role in bottom-up emotional appraisal.^58^ The presence of such adult-like subcortical synchronization at birth suggests that the network is evolutionarily optimized to ensure early survival.^19^ Connectivity between the amygdala and the cortical salience network nodes develops over the first year of life and is characterized by an increase in connectivity over time, with infants reaching adult-level connectivity by 12 months.^19^ Our work tracing the development of anterior insula – amygdala connectivity from fetal to neonatal periods suggests that the two nodes show positive coactivation already at 31 weeks PMA with no major changes during the fetal to neonatal transition. Therefore, the insula-amygdala connection may represent an early-maturing experience-expectant functional infrastructure that prepares newborns to select, evaluate, and integrate sensory information and to learn and adapt to their environment. Given the early interconnectedness between these major cortical hubs and their support of information integration that forms the foundation for numerous complex cognitive functions, their perturbation may have long ranging developmental consequences.

The significance of our findings in the context of autism lies in the presumed role that the communication between insula and amygdala plays in human bonding. A recent model of social development proposes that newborns learn to associate interoceptive signals related to their allostatic needs (hunger, temperature, pain) with exteroceptive signals related to caretaker actions aimed at regulation of the newborn’s allostasis (feeding, warming, soothing) into a multimodal experience.^20^ The inability of newborns to maintain their own allostasis primes them to learn rapidly what does maintain it, which, in the case of human babies, are parents or caregivers. The learning process through which babies come to associate their allostatic regulation with a caregiver provides a foundation for social motivation. Importantly, neural systems that support social behaviors overlap with neural systems supporting allostasis, including insula and amygdala circuitry. Alterations in communication between the two key nodes may contribute to weakening the very mechanism that supports development of social engagement and motivation. Consistent with this view, our findings show significant links between attenuated anterior insula – amygdala connectivity and later social-communication vulnerabilities. Notably, functional connectivity as measured during the first postnatal months reflects both genetic predispositions as well as post-natal experiences within the newborn’s native environment.^59^ Thus, it is important that the associations observed in our study remained robust after maternal stress and mental health indices were considered in the analytic model. The study identifies left lateralized anterior insula - amygdala connectivity as a potential target of further investigation into neural circuitry that enhances likelihood of future onset of behavioral symptoms in autism.

Interestingly, more pronounced hypoconnectivity was observed in male siblings of children with autism compared both to LL female and LL male newborns, with HL females largely aligned with the LL participants. Although due to the very small sample size the results must be interpreted with caution, they are consistent with the notion that male siblings of children with autism are more likely to develop symptoms of autism compared to female siblings^60^ and that in females in general, autism symptoms may arise from different genetic and epigenetic factors and that the likelihood of developing autism symptoms in females is attenuated due to putative protective biological or cognitive factors.^61^ The present study suggests that the sex differences previously described in infant siblings on a behavioral level^62–64^ may also be reflected in brain connectivity patterns in neonates, and motivates further studies into sex differences in brain development both in children with autism as well as their non-autistic first-degree relatives.

Amongst the strengths of the study is a very stringent methodological approach to FC MRI data acquisition and processing as indexed by low FTF values and high total duration of the available time series for each neonate. Further, considering multifactorial contribution to the developing connectome and behavioral outcomes in infancy, accounting for maternal mental health status in all analytic models represents another important area of strength. While differences in FC indices between HL and LL neonates have been reported previously, lack of phenotypic follow-up data has limited interpretation of some of the findings. Here we demonstrate, for the first time, that insula – amygdala hypoconnectivity is linked to the presence of social communication vulnerability in the second year of life. Unique to this study is an inclusion of a fetal sample and our ability to demonstrate using a prospective longitudinal design that the insula and amygdala coactivate already at 31 weeks, providing functional infrastructure for the experience-dependent postnatal development. Interpretation of the findings may be limited by the small HL sample size and a relatively small proportion of participants with 2-year diagnostic outcomes. Larger samples may enable us better to address the question of sex-linked FC effects in infants with familial history of autism, and to detect other alterations in resting state functional connectivity in neonates, contributing to a more complete picture of the impact of genetic predispositions toward autism on early brain development. It will be crucial to replicate the findings in a larger cohort and to extend it into the first months of life to track developmental dynamics of the observed brain-based vulnerabilities and their contribution to behavioral outcomes in infants with familial history of autism.

## Conclusions

The study suggests that the circuitry heavily involved in early development of social boding and motivation may be hypo-connected in those with genetic predisposition for autism by 4 postnatal weeks and that this hypoconnectivity is linked with later emerging social vulnerabilities. Considering that the anterior insula – amygdala axis may be maturing rapidly during the last trimester of pregnancy, our findings motivate future studies into the development of the insula and amygdala networks during the key transition from pre-to postnatal environment and their contribution to later behavioral outcomes relevant to autism.

## Author Contributions

KC, TC, DS, JC, and LM developed the initial aims and design of the study. SM, AV, RF, and AB contributed to study implementation and data acquisition. EB-W managed data sets and contributed to statistical analysis. DS and CL were responsible for MRI data processing and analysis, JC, DS, and KC conducted statistical analysis. KC wrote the manuscript and all authors provided comments and contributed to the revisions.

## Additional Contributions

We thank the children and their families for participating in the study. We acknowledge the clinical team of the Yale Social and Affective Neuroscience of Autism Program including Kelly Powell, Chelsea Morgan, Megan Lyons, and Amy Carney, for their contribution to sample characterization and data collection.

## Author Information

Authors do not have any competing financial interests to disclose.

