## Supplemental Figure 1 for "Hypoconnectivity between anterior insula and amygdala in neonates with familial history of autism"

**Supplemental Materials**

**Figure S1:** Regions of interest used for seed connectivity. The right and left insula seeds are shown in orange and red, respectively.

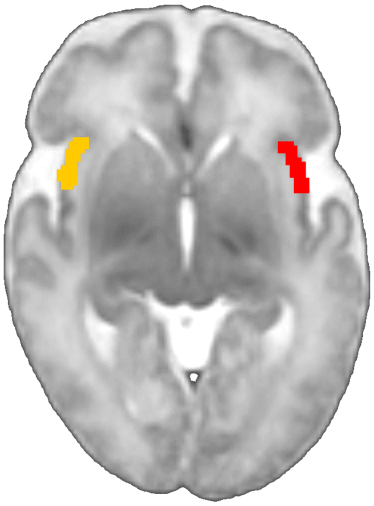
